# Quantifying circulation and antibody cross-reactivity for co-circulating flaviviruses: the case of Usutu and West Nile virus in blackbirds

**DOI:** 10.64898/2026.05.28.728479

**Authors:** Mariken M. de Wit, Nathanaël Hozé, Mart C.M. de Jong, Marion Koopmans, Tjomme van Mastrigt, Reina S. Sikkema, Quirine A. ten Bosch

## Abstract

Serological testing is important for assessing past exposure and immunity, but interpretation can be complicated by antibody cross-reactivity between closely related viruses. We assess this challenge for Usutu virus (USUV) and West Nile virus (WNV), flaviviruses that recently emerged in Europe.

We analysed samples from wild blackbirds collected in the Netherlands between 2016-2022. Samples (N=1742) were screened using an NS1-based protein micro-array, with positives confirmed by Focus Reduction Neutralization Tests (FRNT). We jointly estimated seroprevalence and antibody responses by fitting a Bayesian latent-variable model to FRNT values. Estimates of homologous and cross-reactive antibody responses were used to improve interpretation of observed titres for serosurveillance.

Estimated seroprevalence varied across time and regions between 4.9% (95%CrI 3.5-6.7) to 18.5% (95%CrI 14.9-22.7) for USUV and between 2.4% (95%CrI 1.3-3.8) to 6.4% (95%CrI 3.9-9.6) for WNV.

These were 1.5 (USUV) to 2.4 times (WNV) higher than estimates based on the current threshold-based algorithm. USUV induced a higher antibody response and was more likely to induce a cross-reactive response than WNV. Our classification algorithm informed by these estimates showed high sensitivity (WNV: 0.88, USUV: 0.97) and specificity (both: >0.99).

Our results illustrate how quantitative frameworks can improve serological interpretation in settings with co-circulating pathogens.

## Introduction

Many mosquito-borne diseases circulate between mosquitoes and animals, with spillover occurring to the human population [1]. Surveillance in animals has proven to be a powerful tool for early detection and monitoring of zoonotic pathogens [2], thereby enabling interventions to reduce the risk of zoonotic infections [3,4]. For example, bird surveillance detected West Nile virus (WNV) in the Netherlands [5] leading to retrospective screening which identified human infections [6]. Similarly, an animal mortality network in Central Africa identified Ebola virus circulation in wild animals prior to human cases [7], allowing for an alert to be sent to the health authorities [7]. Beyond early warning, animal surveillance provides a means to monitor spatio-temporal trends in circulation of zoonotic pathogens, as illustrated by wild bird surveillance for Usutu virus (USUV) in the Netherlands [8,9], dead bird monitoring for WNV in the United States of America [10] and livestock surveillance for Rift Valley Fever virus in Uganda [11].

A central pillar of surveillance is serological testing in a target population. By detecting virus-specific antibodies, serological surveillance enables estimation of past exposure, assessment of population immunity, and monitoring of circulation trends in space and time [12]. However, the interpretation of serological assays, including seroneutralisation assays, can be complicated by antibody cross-reactivity of closely related viruses [13,14]. Antibodies generated upon exposure to one virus may bind to antigens of another virus, leading to biased estimates of seroprevalence and misclassification of past infections. This is becoming an increasingly pressing issue as mosquito-borne diseases, for example of the family *Flaviviridae*, expand their geographical ranges and co-circulation becomes more common. This expansion has been observed across the world, as illustrated by the spread of WNV in Europe [15] and emergence of Zika virus in the Amazon region [16] both of which lead to antibody responses that can cross-react with dengue antibodies [13,17].

Another example of emerging co-circulating flaviviruses are USUV and WNV. These viruses have recently emerged and caused outbreaks in Europe [15,18]. While WNV has already been observed in Europe for decades, the number of regions reporting human WNV cases is increasing [15,19]. Both viruses are transmitted in a cycle between *Culex pipiens* mosquitoes and a wide range of bird species [20]. USUV emergence in Europe has led to large-scale mortality in Eurasian Blackbirds (*Turdus merula*, hereafter: blackbird) [21–24]. It has also been detected in humans [25,26], but the impact of USUV on human health remains limited. For WNV, over 3500 human cases have been reported since 2012 in EU/EEA countries (population around 450 million) [19]. In the Netherlands, USUV was first detected in April 2016, followed by WNV in 2020 [8]. Although few WNV-positive birds have been detected since, serological evidence suggests continued circulation [8]. Both viruses belong to the *Flaviviridae* family and the Japanese encephalitis serocomplex [27,28], sharing antigens that make cross-reactive antibody responses common. With the circulation of both viruses expected to increase in future scenarios [29], disentangling potential cross-reactivity between USUV and WNV antibodies becomes increasingly important to ensure effective serological surveillance and reliable patient diagnostics.

Here, we addressed this gap by combining long-term serological surveillance with modelling. Using multi-year blackbird surveillance data from the Netherlands (2016-2022), we developed a Bayesian framework that explicitly accounts for homologous and cross-reactive antibody responses and estimates antibody response parameters for both WNV and USUV. This allows us to propose an alternative estimate of the prevalence of historic infection. Moreover, we account for USUV’s high infection mortality in blackbirds to estimate the attack rate by region and time period. Building on these estimates, we propose and evaluate an individual-based classification algorithm to estimate a bird’s exposure history to improve the utility of serology in surveillance.

## Methods

### Data collection and preparation

Live free-ranging blackbirds were caught and sampled by trained volunteer ornithologists at different locations across the Netherlands from March 2016 to December 2022 (Supplementary figure 1). Sampling was performed under ethical permits AVD801002015342 and AVD80100202114410 issued to NIOO-KNAW. Blackbirds were captured using mist nets and ringed. Blood samples were taken and the date and location of sampling were noted, as described previously [8].

A total of 2051 blackbird sera with recorded sampling location and date were collected. Of these, 309 samples were removed from the analyses because they had low blood volume and hence could not be fully tested or were tested on a different neutralisation test. The remaining 1742 sera were tested for USUV and WNV antibodies in a two-step algorithm as described in [8]. Samples were first screened using a multiplex NS1-based protein micro-array (PMA). Second, samples showing a signal above background (those with a response ≥ 6000 relative fluorescence units (RFU) when tested in a 1:80 dilution) were tested for viral neutralization using Focus Reduction Neutralization Tests (FRNT) (N=461), of which 414 tested positive. In addition, 47 PMA negative samples were included for validation purposes. Sera with a PMA response < 6000 RFU and those with a neutralizing titre < 1:40 to either virus were considered negative for our analysis and their values were set to 0. The use of an alternative cut-off value (< 20) was explored in a sensitivity analysis.

The study period was divided into two periods. Period 1 ranged from March 2016 to April 2020, capturing the years before the first WNV detection by RT-PCR in the Netherlands [5], while period 2 ranged from May 2020 to December 2022. Rather than using calendar years, we split the periods in May to align better with the start of the transmission season. We also divided the country into a Southern (region 1) and Northern (region 2) region. This division was based on the median latitude in the dataset, such that the data were equally distributed across both regions.

### Overview of analyses

In this study, we combine seven years of serological data from wild birds with a Bayesian modelling framework to disentangle antibody cross-reactivity and infer past infection events of USUV and WNV. We analyse virus neutralisation titres obtained through a serological assay and use these data to jointly estimate antibody responses and past exposure probabilities (Figure 1). To facilitate interpretation, we distinguish between three related but conceptually different quantities:

**Figure 1.**
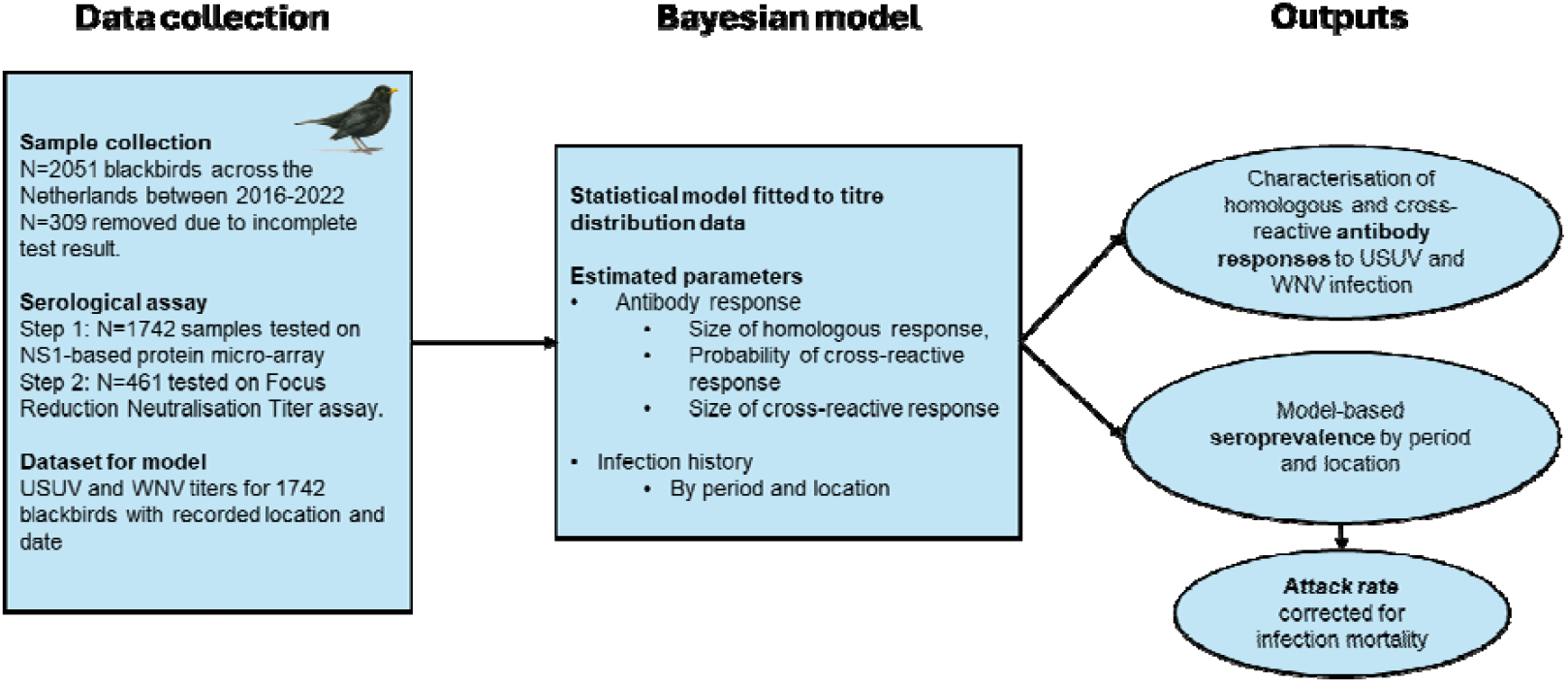
Overview of data collection, model, and outputs. Blackbird samples were collected and analysed through a two-step serological assay (protein microarray followed by FRNT). A Bayesian model was developed to jointly estimate the infection history and antibody responses, accounting for cross-reactivity. This results in estimates of the seroprevalence, the attack rate, and an individual infection history based on the serology.

- Serological data: individual bird samples with a specific combination of USUV and WNV antibody titres.
- Model-based seroprevalence: probability of past infection as estimated by the model. This is based on titres following from homologous and cross-reactive antibody responses after infection. This seroprevalence is estimated by region and time period.
- Attack rate: cumulative probability of infection in the population. This is estimated based on the model-based seroprevalence, corrected for infection-induced mortality. Attack rates are estimated by region and time period.

Together, these definitions provide a framework to interpret serological data and clarify how different analytical approaches relate to underlying epidemiological processes.

### Statistical model

We model the WNV and USUV titre distribution across time and regions, by adapting a model developed for Chikungunya and O’nyong nyong virus in Mali [30]. The model jointly estimates the probability that a host has been infected and the antibody response upon infection. More precisely, because the infection status of an individual is unknown, the model evaluates the probability of a titre response for a given infection scenario weighted by the probability of this scenario.

The likelihood of observing a specific combination of WNV and USUV titres in an individual *j* is given by

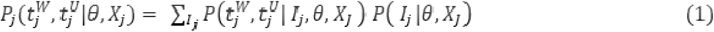

Here, *I*_*j*_ is the history of infection by WNV and/or USUV respectively, resulting in four possible values: 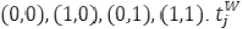and 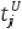 are the normalized ^2^log titres of WNV (*W*) and USUV (*U*), following the formula 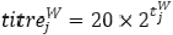 for 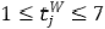 and 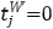 orresponding to any titre below 40. Similarly for USUV, 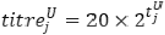 for 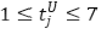 and 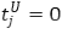 for any titre below 40. We call *P(I*|*θ*, X_*j*_,) the infection model and 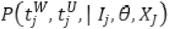 the antibody response model. These probabilities depend on a vector *X*_*j*_ of individual covariates (region and period) and the parameter vector *θ = (σ*^*w*^,*σ*^*U*^,*σ*^*w*→*U*^,*σ*^*U*→*W*^*p*^*W*→*U*^,*p*^*U*→*W*^,*λ*^*W*^,*λ*^*U*^,*β*^*W,per*^,*β*^*U,per*^,*β*^*W,loc*^,*β*^*U,loc*^), which are all the parameters for the antibody response and the probability of infection. These parameters are more specifically estimated in the *antibody response model* and the *infection model*, respectively, which are described below.

#### The antibody response model

The antibody response depends on the history of infection. We assume the titres generated by a homologous infection or cross-reactivity are stochastic and non-zero. The model accounts for distinct response parameters for USUV, WNV, and the individual cross-reactive responses. Additionally, the probability of a cross-reactive response is estimated as part of the model.

Formally, in the absence of infection, both normalised log titres are 0 and

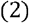

where *δ*(*t*) is 1 if t=0, and 0 otherwise. If a bird is infected by WNV but not by USUV, then we assume a positive homologous response for WNV, and a cross-reactive USUV response may arise with a certain probability. The joint probability of observing titres is composed of three components: the first term characterises the homologous response parameterized by σ^W^. The distribution of the response *f* is a zero-truncated Poisson distribution. The second and third part concerns the cross-reactivity. When only the titre against one virus was positive, no cross-reactivity occurred. A cross reactive response therefore happens with probability *p*^*W*→*U*^ and is of size *σ*^*W*→*U*^ on average, which is lower than the homologous response such that *σ*^*W*→*U*^*=*^*W*→*U*^*\*σ*^*U*^,where ^*W*→*U*^≤1. As a consequence there is no cross-reactivity with a probability 1−*p*^*W* →*U*^ Overall the probability of observing the titres is

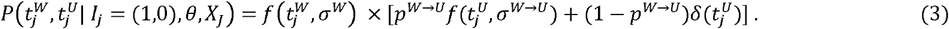

Similarly if there was an infection by USUV but not by WNV, then

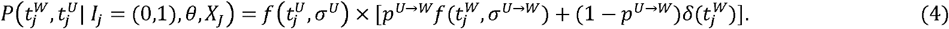

When the bird has been infected by both viruses, the titre probability distribution is a weighted sum of the probability of cross-reactivity,

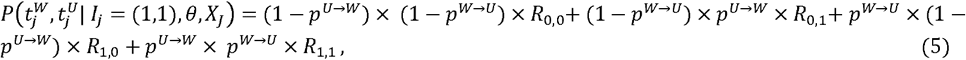

where, for convenience, *R*_0,0_ denoted the response in the absence of cross-reactive response, *R*_0_,_1_,is the response when WNV induces a cross-reactive response and USUV does not, and so on. They are derived from equations (3) and (4) and can be expressed as a convolution, meaning that the observed response is the sum of homologous and cross-reactive components. The four response terms are

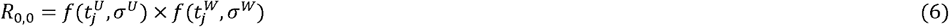

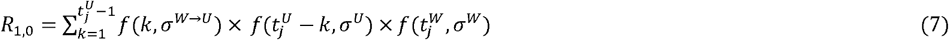

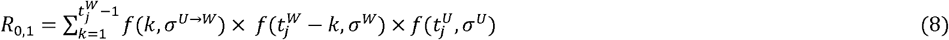

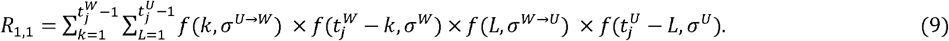

#### The infection model

We consider that the probabilities of infection by WNV and USUV are independent, and may vary by sampling period and location. The probability of infection by WNV and USUV is

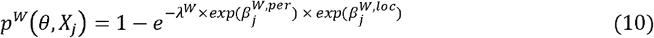

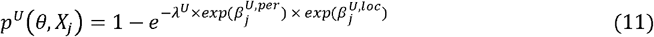

Where 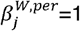 in period 1 (years 2016-2020) and 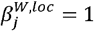in location 1 (South). The parameter λ^*W*^ is a reparametrization of the probability of infection of WNV in period 1 and location 1. For ease of use we will call force of infection (FOI) the λ parameters, although they characterize a probability for a sample to be positive, and not the rate of infection over a fixed time interval. For instance, a positive bird in period 2 may have been infected during period 1.

The infection model characterizes the probability of each of the four scenarios of infection.

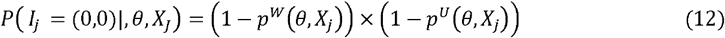

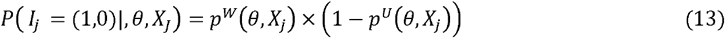

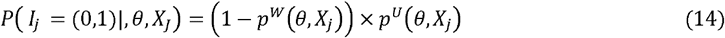

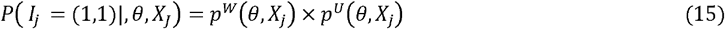

#### Implementation

The model was fitted to the serology data using a Markov chain Monte Carlo (MCMC) framework through the *rstan* package [31]. We used a No-U-Turn sampler to sample parameter values. We ran three chains of 2000 iterations using a burn-in of 1000 iterations. All prior distributions were normally distributed. We used mean 0 and standard deviation 3 for the period and location effects 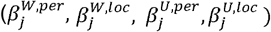,for the homologous antibody responses (*σ*^*W*^, *σ*^*U*^), and for the FOIs (λ^*W*^, λ^*U*^). Prior distributions for the probability of a cross-reactive response (*p*^*W*→*U*^,*p*^*U*→*W*^) had a mean of 0 and a standard deviation of 1, while for the size of the cross-reactive response (*σ*^*W*→*U*^, *σ*^*U*→*W*^)the mean was 0 with a standard deviation of 0.25. Convergence was assessed using standard tools in rstan (ESS, Rhat, number of divergences) and traceplots (Supplementary Figure 3).

### Model outcomes

The model jointly estimates parameters related to the homologous and cross-reactive antibody responses as well as parameters related to the infection probability (i.e. corrected seroprevalence). While the parameters related to the antibody responses can be interpreted directly, additional adjustments are required for the infection probability-related parameters due to infection mortality.

### Adjustment for mortality

Because the mortality rate in blackbirds due to USUV infection is high (~70%) [9,21,24], the measured USUV seroprevalence underestimates the true attack rate of the virus. To correct for the bias introduced by this high mortality, we model the infection of a population of *N* susceptible birds and

Write *q*^*U*^ the probability of infection by USUV during a period (the attack rate) and *δ*^*U*^ the mortality rate. At the end of this period, *N*(1− *q*^*U*^) remain uninfected, *Nq*^*U*^*δ*^*U*^ have died and *Nq*^*U*^ (1− *δ*^*U*^) have survived. The observed seroprevalence is therefore

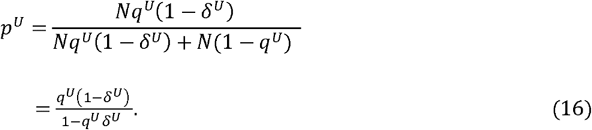

The attack rate is retrieved from the observed seroprevalence with the formula

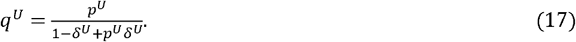

In Supplementary Figure 7A, we represent the relationship between the true attack rate and seroprevalence for various values of the infection mortality rate. The value for infection mortality risk used in our analysis was obtained from de Wit et al. [9] (median 74%, 95%CI 66-78). This correction assumes that natural mortality is independent of infection status. Under this assumption, background mortality affects infected and uninfected individuals equally and therefore cancels out when computing seroprevalence among surviving individuals. In addition, we assumed a closed population over the observation period, or equivalently that population turnover does not alter the infection status distribution of sampled individuals. In practice, recruitment of naïve individuals or changes in population size may further dilute seroprevalence and lead to additional underestimation of the true attack rate.

### Model evaluation

We evaluated the ability of our model to estimate parameters from simulated data. We created three simulated datasets using three different sets of parameters. The number of individuals in each simulated dataset was equal to that in the real dataset (N=1742). For each simulated dataset, we evaluated the median and 95% credible interval of the posterior distributions and compared this to the input parameters (see Supplementary figure 2).

Additionally, we evaluated the quality of the model fit to the real data by simulating 1000 datasets (of 1742 observations) with parameters drawn from the posterior distributions. We compared the observed number of samples with given WNV and USUV titre values per period and location with the simulated number.

Our model provides an estimate of the seroprevalence, accounting for antibody cross-reactivity. Currently, other classification algorithms are used to assign the serostatus based on observed titres, resulting in different estimates of the seroprevalence. We assessed whether the model-based seroprevalence provides a better representation of the observed titre distribution compared to the seroprevalence estimated using the current algorithm (Supplementary figure 6, see “serological classification” for a description of the current algorithm). We refitted the model to the observed data while fixing the seroprevalence at values estimated from the current classification algorithm, thereby only estimating the antibody response parameters. We then compared the quality of the model fits to the real data between these two models.

### Sensitivity analyses

To explore the impact of several assumptions on the results, we compared three model versions in a sensitivity analysis. In model S1 we assumed no WNV circulation prior to its detection in 2020. This was implemented by setting the FOI in period 1 to zero and removing the parameter 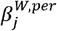 such that for period 2

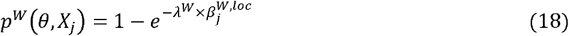

In model S2 we assumed no cross-reactive antibody responses. This was implemented by removing the probability and size of the cross-reactive response from the model. In model S3, we assumed all titre values ≥ 20 were positive, leading to an additional level for *t* now ranging from 0 to 8. Models were compared using the leave-one-out information criterion (LOOIC).

### Serological classification

We used the model to develop and evaluate a classification algorithm based on the titre combination and the estimated seroprevalence of both viruses. The algorithm was developed at the estimated seroprevalence level and evaluated under alternative seroprevalence scenarios.

First, we simulated the distribution of titres after an infection with USUV, WNV, or both, using the posterior distributions from the antibody response model. Second, we calculated the probability of each titre combination in the population by weighting these distributions by the estimated seroprevalence. Then, we inverted this relation to estimate the probability that an individual has been infected by USUV, WNV, or both, given their titres. Each titre combination was assigned to the most likely infection status. Finally, we calculated sensitivity and specificity of this classification rule. For WNV (analogously for USUV), the true and false, positive and negative rates were obtained by the weighted sum of the relevant titre combinations multiplied by the probability that the titre was or was not attributable to an infection.

We compared this model-based classification rule to the currently used algorithm for classification (i.e. threshold-based classification), which was developed based on confirmed positive and negative patient and animal host sera. In the threshold-based classification algorithm, a sample is considered USUV positive when PMA ≥ 6000, USUV FRNT ≥ 160, and USUV FRNT ≥ 4* WNV FRNT. A sample is considered WNV positive when PMA ≥ 6000, WNV FRNT ≥ 80, and WNV FRNT ≥ 4* USUV FRNT. Samples with PMA ≥ 6000 and an FRNT value above the threshold, but not 4 times higher on one virus, were considered ambiguous. Samples with no reactivity on PMA or an FRNT value below the threshold were classified as negative.

## Results

### Serology of USUV and WNV

Blackbird blood samples were collected and tested in several locations across the country between 2016-2022 (Figure 2C). Samples were included in the analysis when their location and date were recorded and they satisfied the full testing algorithm, resulting in a total of 1742 samples from 1496 individual birds. The proportion of samples positive on the PMA (step 1 of the testing algorithm) varied by year and region (Figure 2A) as did the FRNT results (step 2 of the testing algorithm) (Figure 2B). The overall proportion of samples positive on the PMA was similar in the years 2016-2019 (25.0%, 95%CI 22.0-28.2) compared to 2020–2022 (22.7%, 95%CI 20.1-25.5), but higher in the South (27.2%, 95%CI 24.1-30.4) compared to the North (20.8%, 95%CI 18.3-23.6). However, the geometric mean FRNT values were higher for USUV (64.5 (95%CI 55.6-74.8) in the first period, compared to 41.2 (95%CI 36.6-46.5) in the second period. The trend was similar for WNV: 27.2 (95%CI 24.9-29.7) in the first period and 21.7 (95%CI 20.2-23.3) in the second period. The regional differences were stronger, with a higher geometric mean titre of USUV in the South (83.3 (95%CI 72.0-96.5), compared to the North (30.9 (95%CI 27.6-34.6)) A similar trend was observed for WNV (South 26.4 (95%CI 24.3-28.8), North 22.2 (95%CI 20.7-23.9)). Overall, regional and temporal trends were similar for both viruses.

**Figure 2:**
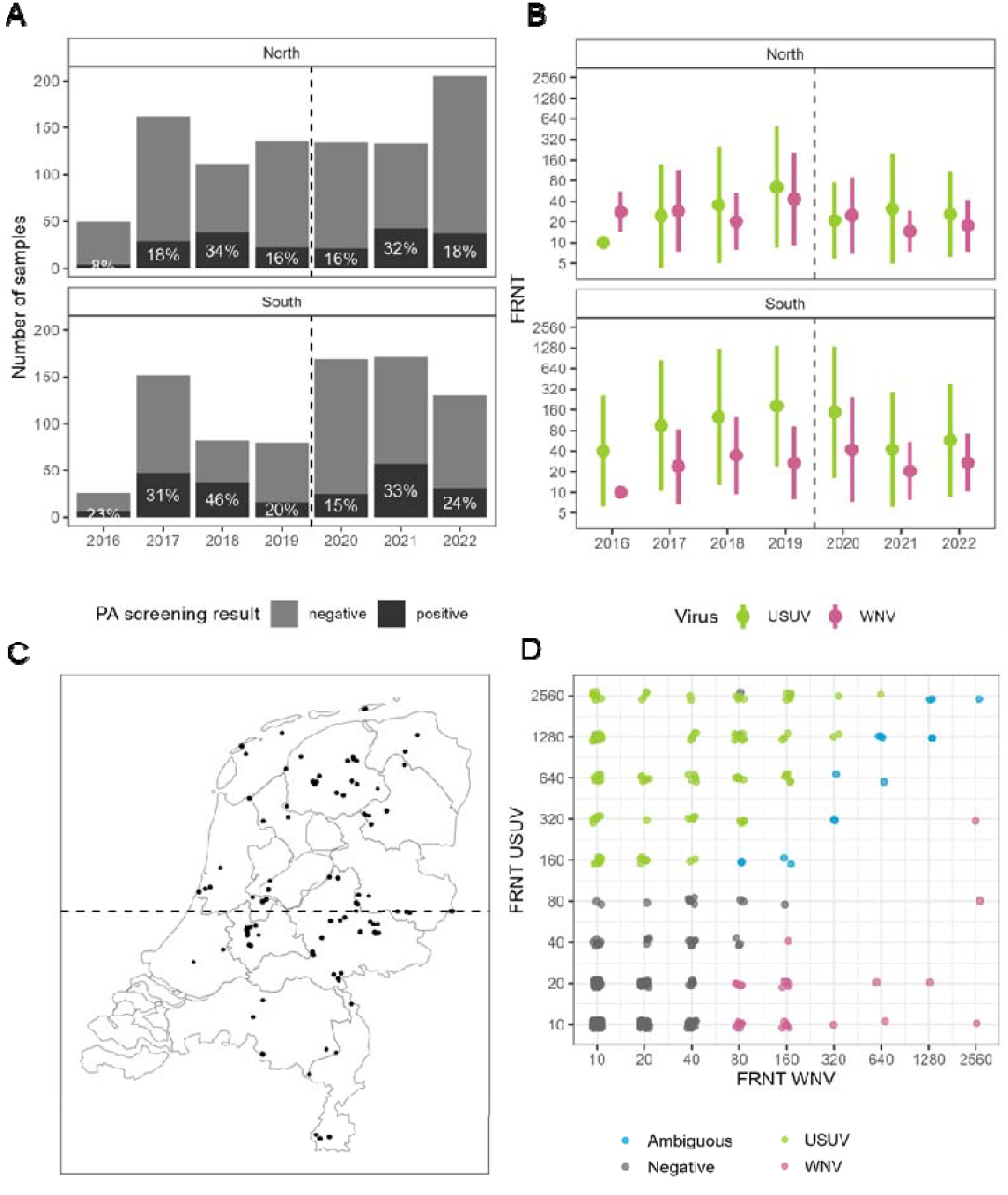
Descriptive results of tested blackbirds. A. Number of samples and proportion PA-positive by year and region. The vertical line indicates the division between period 1 and 2. B. Distribution of FRNT values (mean and standard deviation) for USUV (green) and WNV (pink) by year and region. C. Map of sampling locations, where each dot represents one sample. The horizontal line indicates the division between North and South. D. Paired FRNT values for USUV and WNV coloured according to their classification status using the threshold-based algorithm.

However, interpretation of the mean FRNT (Figure 2B) is difficult. First, the titres span several orders of magnitude and the variation is larger for USUV than for WNV, making the comparison of the titres for the two viruses hard to interpret. Second, the mean FRNT includes samples from infected and non-infected birds, so a high mean FRNT could indicate a strong antibody response in some birds or a high overall prevalence. Third, potential cross-reactivity further complicates the interpretation. The threshold-based algorithm finds that 7.5% of samples were USUV-positive, 1.7% were WNV-positive, and 0.9% were ambiguous (Figure 2D). The ambiguous samples are assumed to be cross-reactive positive samples or dual positive, with FRNT titres less than 4 times higher between the viruses.

### Disentangling antibody responses

To provide an interpretation of the observed titre combination, we developed a mathematical model that explicitly describes the antibody response of WNV and USUV in blackbirds, accounting for cross-reactivity between the two viruses. The model jointly estimates the homologous and cross-reactive antibody responses to an infection together with the prevalence of historic infection (called “seroprevalence” throughout) by period, location, and virus.

We found that the homologous titre response to infection was larger for USUV (geometric mean FRNT 509 (95% credible interval (CrI) 406-636)) than for WNV (101 (95%CrI 85-122)) (Figure 3). Moreover, USUV was more likely to induce a WNV cross-reactive response (in 50.9% of infections (95% CrI 43.0-58.3) compared to 12.2% (95%CrI 1.4-25.6) vice versa). The cross-reactive response of an USUV infection on WNV antibody titres (88 (95%CrI 75-105)) tended to be larger than the cross-reactive response of an WNV infection on USUV antibody titres (53 (95%CrI 40-110)), but posterior distributions overlapped. To test if our model adequately represented the data we then simulated serological assays based on posterior parameters and found that our model fit very well to the observed data, with 97% of observed values falling in the range of simulated values (Supplementary Figure 5). We performed several sensitivity analyses. We changed the positivity cut-off for titre values, which showed that model fit was worse when titre values ≥ 20 were treated as positive (LOOIC 4273 compared to 3129 in the main model when only values ≥ 40 were positive). We also removed the possibility of cross-reactivity from the model, which significantly worsened the model fit (LOOIC 3439 compared to 3129 in the main model with cross-reactivity).

**Figure 3:**
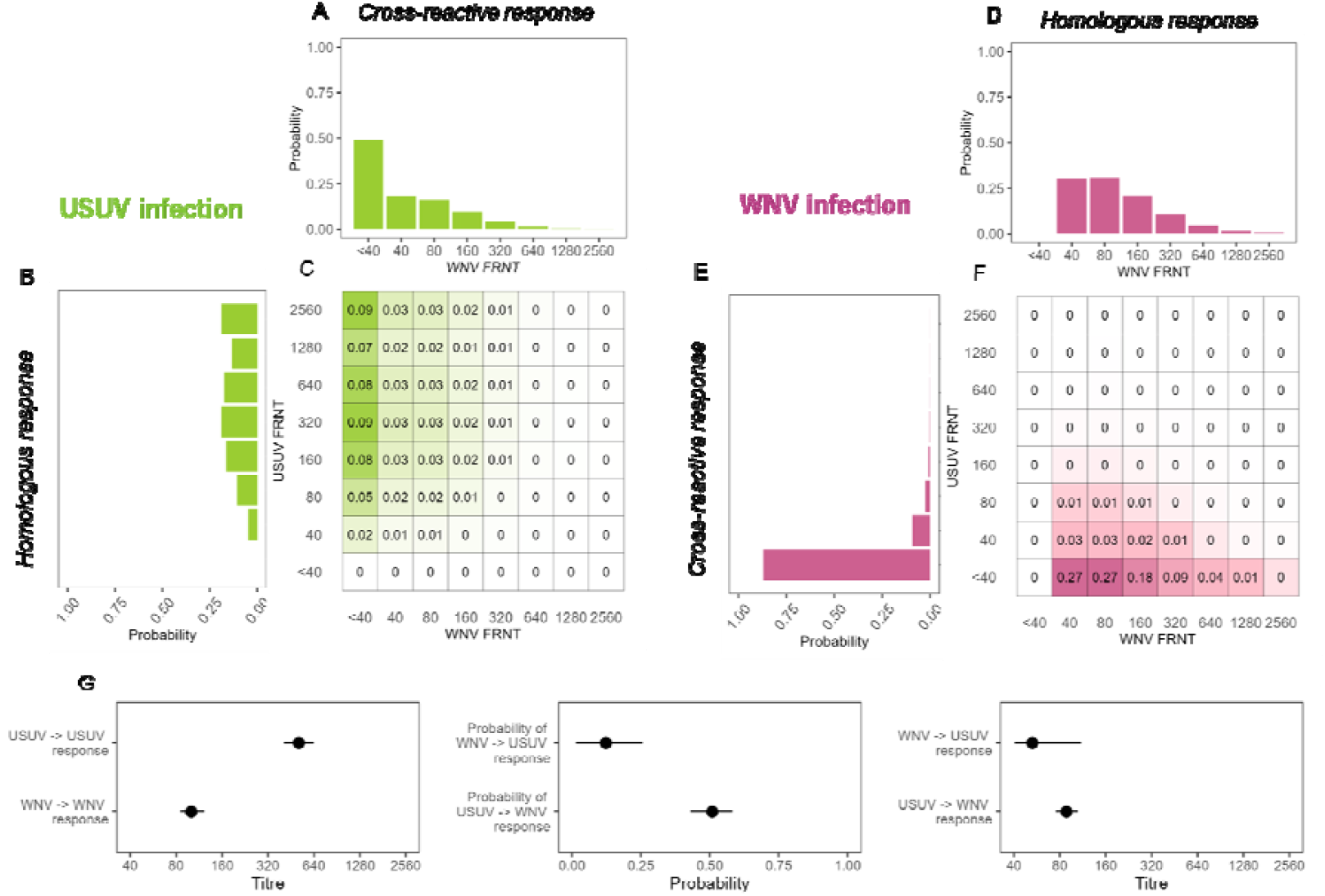
Characterisation of antibody response to infection. A-F. Model estimated distribution of antibody titres to infection with USUV (a-c, in green) or WNV (d-f, in pink). Heatmaps show the joint distribution of USUV and WNV FRNT values following infection. Histograms are the sums along rows and columns and show the homologous responses (b, d) and cross-reactive responses (a, e). G. Mean and 95%Crl of posterior distributions related to the antibody responses. The first panel shows the size of the homologous antibody responses (σ^U^andσ^w^), the second panel shows the probability of cross-reactivity antibody responses (p^w→U^and p^U→W^), the third panel shows the size of the cross-reactive antibody responses (σ^w→U^and σ^U→W^). See supplementary figure 4

### Estimating infection history

Concurrent with antibody responses, the model estimated the cross-reactivity corrected seroprevalences. Similar to estimates from the threshold-based algorithm, the model estimates also show higher seroprevalence for USUV compared to WNV (Figure 4A). For both viruses, seropositivity was higher pre-2020 compared to 2020-2022 and in the South compared to the North. For WNV, seropositivity varied between 2.4% (95%CrI 1.3-3.8) in the North between 2020-2022 to 6.4% (95%CrI 3.9-9.6) in the South 2016-2020. For USUV, these values were from 4.9% (95%CrI 3.5-6.7) to 18.5% (95%CrI 14.9-22.7). Even though no WNV PCR-positive birds were found prior to 2020, model comparison showed that fit was better when allowing for a non-zero WNV infection probability prior to 2020 (LOOIC 3120 in the main model, compared to 3330 in an alternative model without early WNV transmission). For USUV, our model estimated a larger seroprevalence than the threshold-based algorithm did with the mean being on average 1.5 times higher. On average, the model estimate of WNV seroprevalence was 2.4 times higher than with the threshold-based algorithm. However, spatial and temporal trends were the same between both methods. Comparison of model fit between this model (estimated seroprevalence) and a model where seroprevalences are fixed at the uncorrected values showed that the data is best explained by the model-estimated seroprevalence (Supplementary figure 6).

**Figure 4:**
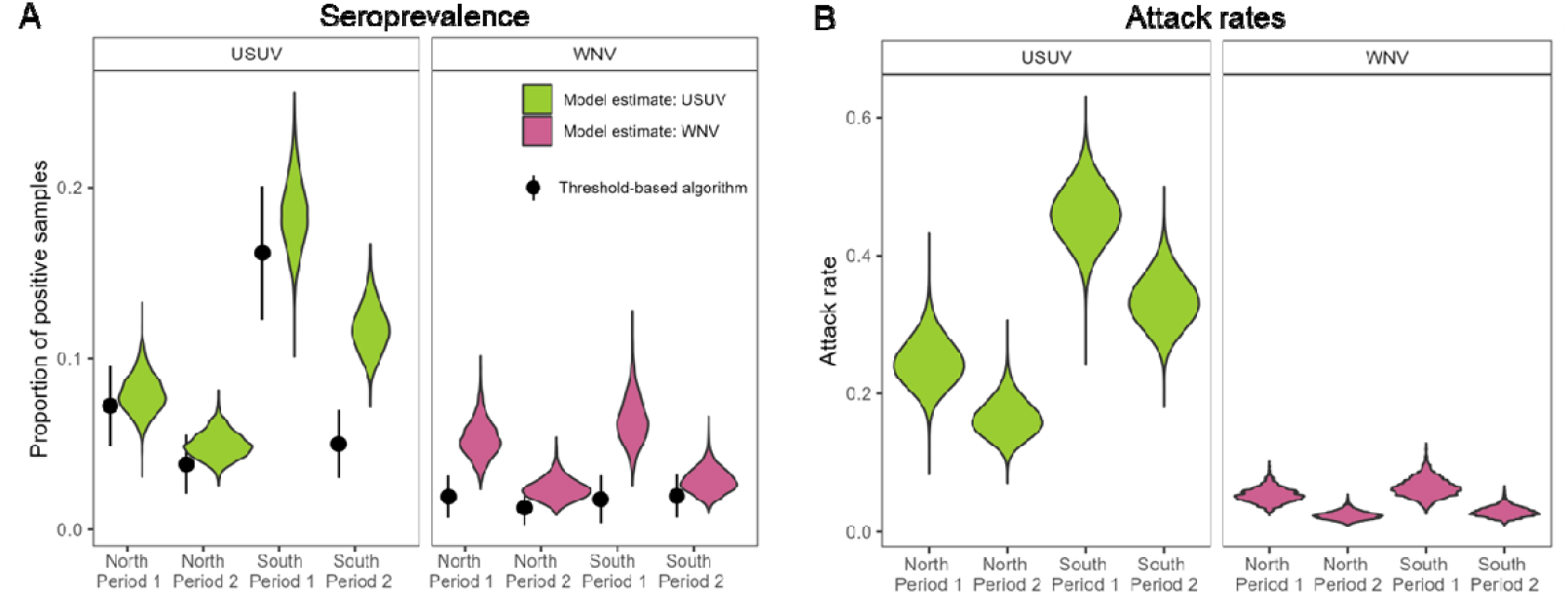
Seropositivity and attack rates. A. Comparison of proportion of samples classified as seropositive using the threshold-based algorithm (black) and the model-based seroprevalence estimates (green for USUV, pink for WNV) split by period and region. B. Estimated attack rates of USUV (green) and WNV (pink), split by region and period. Estimates for USUV are corrected for infection-induced mortality.

To estimate attack rates, we adjusted the model-based seroprevalence for infection-induced mortality of USUV. Estimates of the attack rate followed the same trends, but were much higher for USUV due to elevated infection mortality (Figure 4B). This showed that while the estimated seropositivity was 2.8 times (95%CrI 1.1-6.2) higher for USUV than for WNV, the attack rate for USUV was 8.9 times (95%CI 2.2-24.1) higher than for WNV.

### Interpreting serological surveys

A key objective of our study was to translate model results into a practical classification algorithm. We reconstructed the population titre distribution from our calibrated model. Applying the current threshold-based classification algorithm to these reconstructed data, we estimated sensitivity of this algorithm at 0.63 for WNV and 0.84 for USUV at the model-derived seropositivity levels. Specificity of WNV and USUV was high (0.989 and >0.99 for WNV and USUV, respectively).

To increase the sensitivity of the threshold-based approach, we proposed a probability-based classification grounded in the estimation of the seroprevalence and antibody response distributions. Our approach estimates the probability that a bird was historically infected with USUV and/or WNV for a given titre combination. Because this probability depends on the overall probability of infection, which is a proxy for the seroprevalence, we provide a simple rule calibrated at the levels of seropositivity estimated by the model (10.8% for USUV, 4.2% for WNV) averaged across region and period.

Using the modelled antibody response to infection (Figure 3), we classified birds according to their most probable infection status (Figure 5A). The resulting model-based classification algorithm is simple: because the cross-reactive USUV response is rarely above 80 (Figure 3F), any sample with USUV titre ≥ 80 is classified USUV-positive, any sample with WNV titre ≥ 40 while USUV titre < 80 is classified as WNV-positive. When WNV titre is ≥2560 and USUV titre is ≥80 it is both USUV-positive and WNV-positive. While this classification is based on most probable infection status, other classification algorithms, such as one that maximises sensitivity for one of the viruses, can also be developed and evaluated based on this heatmap (Figure 5A).

**Figure 5:**
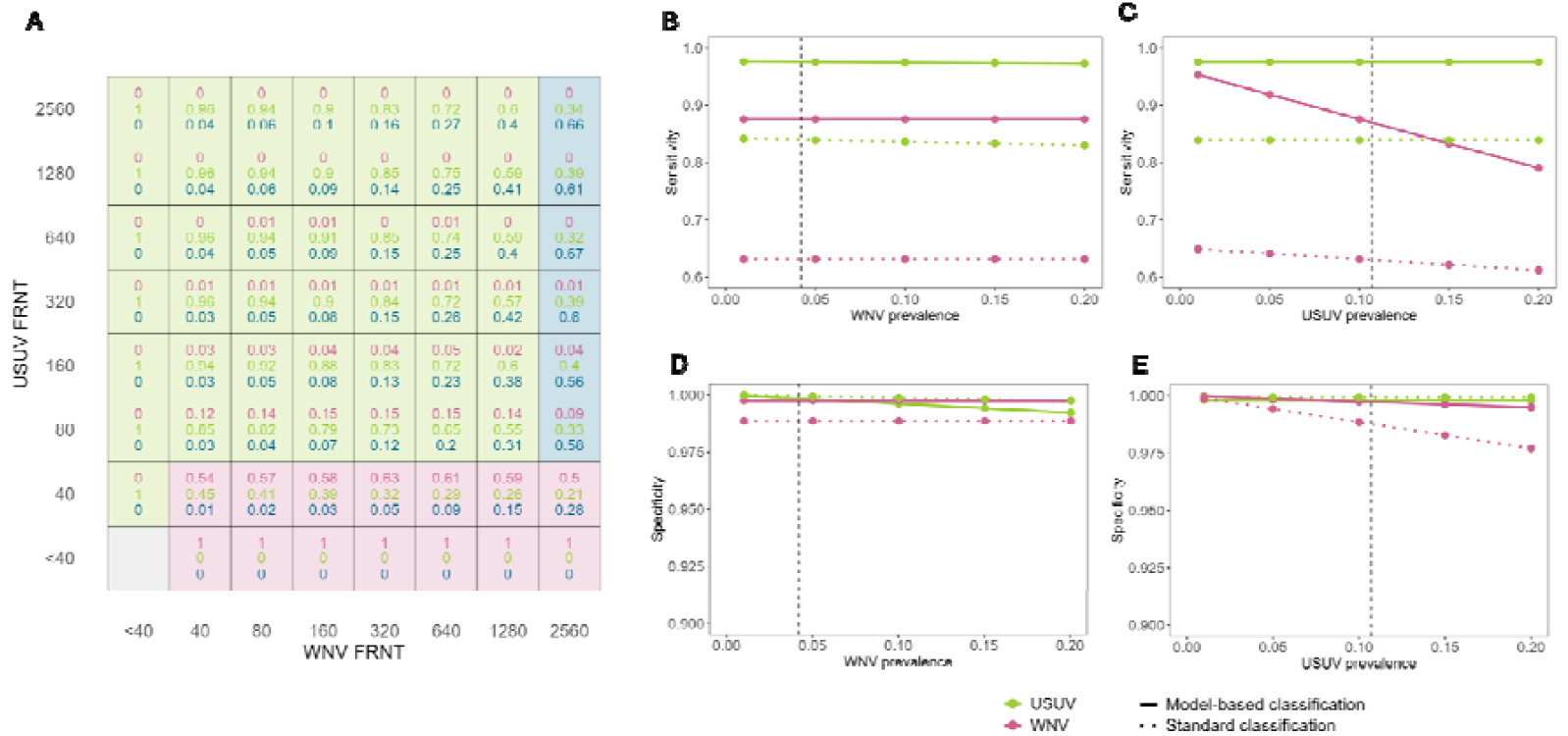
Titre classification algorithm. **A. Historic infection probability heatmap.** Values in cells represent the probability of the observed titre combinations being the result of a historic WNV infection (pink), USUV infection (green), or co-infection (blue). Cells are coloured according to the most probable infection status with WNV infection in pink, USUV infection in green, and co-infection in blue. This heatmap is created for the observed seroprevalence values (10.8% for USUV and 4.2% for WNV) using the estimated antibody response parameters. **B-E. Sensitivity & specificity of proposed classification algorithm**. The classification rule is optimised for estimated current seropositivity levels (10.8% for USUV, 4.2% for WNV). Lines indicate the effect of variation in seroprevalence of one virus on the performance of the algorithm, while the seroprevalence of the other virus is kept constant.

This classification substantially improved performance, with sensitivity increasing to 0.98 for USUV and 0.87 for WNV. Specificity was >0.99 for both viruses for the estimated seropositivity levels. Because we proposed a classification that is optimised for current seroprevalence levels, performance is somewhat reduced for different seroprevalence levels. However, sensitivity and specificity for USUV only decreased slightly when WNV seroprevalence increased (Figure 5B,D). In contrast, higher USUV seroprevalence decreased the sensitivity for WNV detection (Figure 5C) due to high probability of inducing a cross-reactive response (Figure 3G), but effects on the specificity for WNV were minor (Figure 5E).

## Discussion

Interpretation of serological surveillance data can be hampered when related pathogens co-circulate. This makes it difficult to determine whether detected antibodies reflect exposure by one or the other pathogen. By integrating modelling with continuous surveillance data, our study illustrates how analytical frameworks can strengthen serological surveillance in contexts of cross-reactivity and co-circulation. We showed this through a concrete application to two closely related flaviviruses that co-circulate in European wildlife: USUV and WNV. We found that seroprevalence is likely higher than estimated based on current threshold-based algorithms for both USUV and WNV in blackbirds in the Netherlands. The difference was larger for WNV, likely because of a lower sensitivity of the currently used algorithm due to WNV inducing a lower antibody response than USUV. Results also indicated that USUV induced a stronger cross-reactive response. We developed an individual-based classification algorithm based on this detailed characterisation of antibody responses. This showed high sensitivity and specificity for both viruses on data simulated for current estimated seroprevalence levels. Due to USUV inducing a higher antibody response, sensitivity for WNV declined with increasing USUV circulation. Despite this, sensitivity was consistently higher than for the threshold-based algorithm. The other performance measures were robust to changes in circulation.

### Spatiotemporal trends

This analysis revealed similar spatiotemporal trends of exposure to USUV and WNV in Dutch blackbirds, with higher seropositivity for USUV than for WNV. Although the first WNV PCR-positive bird was not found until 2020, we found strong statistical evidence of circulation before 2020. WNV detections have been low throughout the study period, with 8 PCR positive birds in 2020 and 1 in 2022 [8]. Interestingly, these detections were not associated with increased WNV antibody titres. In fact, a lower mean WNV titre was observed in the 2020-2022 period (Figure 2B). This was reflected in our seroprevalence estimates which suggest that circulation of both viruses was lower after 2020 than before. For USUV, this aligns with previously observed high PCR-prevalence before 2020 [8] and blackbird population decline [32] between 2016-2018. It is possible that high USUV circulation levels before 2020 resulted in repeated infections in the same birds, which might result in a different antibody response compared to a single exposure. Additionally, antibody responses after repeat exposure might differ depending on the time since first infection due to original antigenic sin [33]. This could have impacted the accuracy of the seroprevalence estimates for both viruses. The similarity in temporal trends between both viruses may reflect shared ecological drivers as both are transmitted by similar mosquito species [20].

### Cross-reactivity corrected seroprevalence

Our results suggest that seroprevalence is underestimated in the current algorithm. Several factors contributed to the higher estimated seroprevalence from our model. First, the positivity cut-off in the threshold-based algorithm was a minimum titre of 80 for WNV and 160 for USUV, while we considered a titre of at least 40 as positive. Second, the threshold-based algorithm requires the titre for one virus to be at least 4 times higher than for the other virus and therefore assumes symmetry in the cross-reactive response. However, we found statistical support for a high degree of asymmetry in the cross-reactive response and based the required difference in titre on these specific antibody responses. Third, while the threshold-based algorithm is unable to classify those samples with high titres without a clear difference between the viruses, the model does not suffer from this. Especially in the South after 2020, the model showed higher estimated USUV seroprevalence compared to the threshold-based algorithm. This could be due to lower observed titres (Figure 2B) being classified as negative in the current algorithm. This difference in sensitivity may imply that the current algorithm is able to detect recent infections (with higher titres), but is more likely to miss older infections with lower titres. These findings suggest that seropositivity levels are higher than previously thought for both USUV and WNV in blackbirds, which might have implications for analyses based on these data, including the infection-induced mortality of USUV in blackbirds estimated in de Wit et al. [9].

A key limitation of our seroprevalence estimates is related to their interpretation across multi-year periods. The estimated seroprevalence reflects the likelihood that a bird sampled within a given period had been infected at some point up to that time at any location it has been (including outside the Netherlands), rather than the period- and location specific forces of infection. Antibody waning is also challenging the interpretation of the data. It was indirectly accounted for by modelling titre values using a zero-inflated Poisson distribution, which allows for low or undetectable titres among previously infected individuals. However, explicit temporal dynamics of antibody decay were not modelled. Given the relatively short lifespan of blackbirds, substantial long-term waning is unlikely to strongly affect population-level estimates, but it remains possible that some historic infections are misclassified as cross-reactive responses. Longitudinal studies could shed light on long-term antibody dynamics in blackbirds. Additionally, we assumed that antibody responses were independent of previous exposure history meaning that the response to a repeated exposure (with the same or the other virus) is the same as to a first exposure. However, repeated exposure in individual birds is likely rare given the relatively low seroprevalence of both viruses in blackbirds.

### New classification algorithm

For practical surveillance applications, we proposed a simplified, model-informed classification algorithm. Compared with the threshold-based approach, this classification showed improved sensitivity and specificity on simulated datasets representing plausible circulation scenarios. However, these improvements were evaluated within the current modelling framework and therefore lack external validation against a gold standard. As noted in recent work on multiplex assays of flaviviruses [34], the advantage of this approach is that it does not require validated samples that are often difficult to obtain. However, further validation using cohort or longitudinal data would strengthen the evaluation of the classification’s performance. Despite this limitation, the model is based on FRNT data, widely regarded as the best available method for quantifying neutralising antibody responses. Sensitivity analyses further demonstrated the robustness of the proposed classification across a wide range of plausible seroprevalence scenarios, with only limited reductions in WNV sensitivity at high USUV seroprevalence. Specificity for both viruses was largely unaffected across scenarios. These results suggest that, even in the absence of external validation, the model-based classification provides a meaningful improvement over the threshold-based approach. The proposed classification algorithm can also be adapted depending on surveillance goals and desired trade-offs between sensitivity and specificity for each virus and between both viruses, such as when detections are used to trigger interventions. Our results can be used to design and evaluate the performance of these different classification algorithms. The proposed classification is specific to blackbirds and highlights that titre classification algorithms can benefit from incorporating host-specific immune responses and epidemiological context.

### Co-circulation and cross-reactivity

Co-circulation of USUV and WNV has been reported across Europe [20], including in Germany [35], Spain [36], and Italy [37]. In Germany, even co-infections have been observed in dead birds [38]. Cross-reactivity may also provide partial cross-protection. An experimental study in geese showed that geese first infected with USUV and later with WNV showed lower viral loads and less severe histopathological lesions than birds only infected with WNV [39]. This study also observed an increase in WNV antibody titre after the initial USUV infection and a boosting of the USUV antibody titre after subsequent infection with WNV, indicating a cross-reactive response in both directions possibly influenced by original antigenic sin. Experimental infections in magpies showed that viremia and WNV antibody titres after WNV infection did not differ between magpies previously exposed to USUV and those that were not [40]. However, they observed higher survival rates after WNV infection in birds previously exposed to USUV, suggesting a protective effect of previous USUV exposure. Interestingly, while we observed a larger homologous and cross-reactive response to USUV infection compared to WNV in blackbirds, experimental infections in red-legged partridges (*Alectoris rufa*) found the opposite, showing higher homologous and cross-reactive titres following WNV infection [41].

Our finding of strong USUV-induced cross-reactive responses suggests that prior USUV exposure may reduce WNV susceptibility in blackbirds if these cross-reactivity antibodies lead to cross-immunity. It is important to note that the analysis presented here is restricted to blackbirds. The extent to which USUV circulation reduced the impact of WNV emergence at the population level depends on the characteristics of these immune responses in other competent host species, the level of cross-immunity this induces, and on the degree of overlap in hosts that are important for each virus’ transmission. Given the large number of host species susceptible to infection with both viruses and the observations of cross-reactive response in several species, it seems likely that previous USUV circulation influenced WNV emergence.

## Conclusion

This study demonstrates the value of applying analytical methods to continuous surveillance data to improve the interpretation of serological population data in contexts with co-circulating viruses. The extent of antibody cross-reactivity combined with high seropositivity levels we observed suggests that cross-immunity should be considered when interpreting epidemiological trends of USUV and WNV.

## Acknowledgements

We would like to thank all volunteers at the bird ringing stations across the Netherlands for their efforts in catching and sampling birds. We would also like to thank all people involved in the coordination of these surveillance schemes as well as those involved in the laboratory analyses (Felicity Chandler). We are grateful to Henk van der Jeugd and Jurrian van Irsel for the insightful discussions.

## Funding

This publication is part of the project ‘Preparing for Vector-Borne Virus Outbreaks in a Changing World: a One Health Approach’ (NWA.1160.18.210), which is (partly) financed by the Dutch Research Council (NWO).

## Data availability

The serology dataset analysed in this study is available via the BioStudies database at http://www.ebi.ac.uk/biostudies under accession codes S-BSST1867 and is also available via the Pathogens Portal Netherlands at https://www.pathogensportal.nl/arboviruses.html.

## Author contributions

**MdW** : conceptualisation, methodology, software, formal analysis, visualisation, data curation, writing – original draft, writing – reviewing and editing. **NH**: conceptualisation, methodology, software, formal analysis, visualisation, writing – original draft, writing – reviewing and editing. **MdJ** : conceptualisation, methodology, supervision, writing – reviewing and editing. **MK**: funding acquisition, writing - reviewing and editing. **TvM**: data curation, writing - reviewing and editing. **RS**: data curation, writing - reviewing and editing. **QtB** : conceptualisation, methodology, supervision, writing – reviewing and editing.

## Supplementary Materials

### Outline

1. Descriptives
2. Identifiability analysis
3. Diagnostics
4. Posterior distributions and model fit
5. Impact of infection-induced mortality on parameter estimates
6. Titre classification

### Supplement: Descriptives

**Supplementary figure 1.**
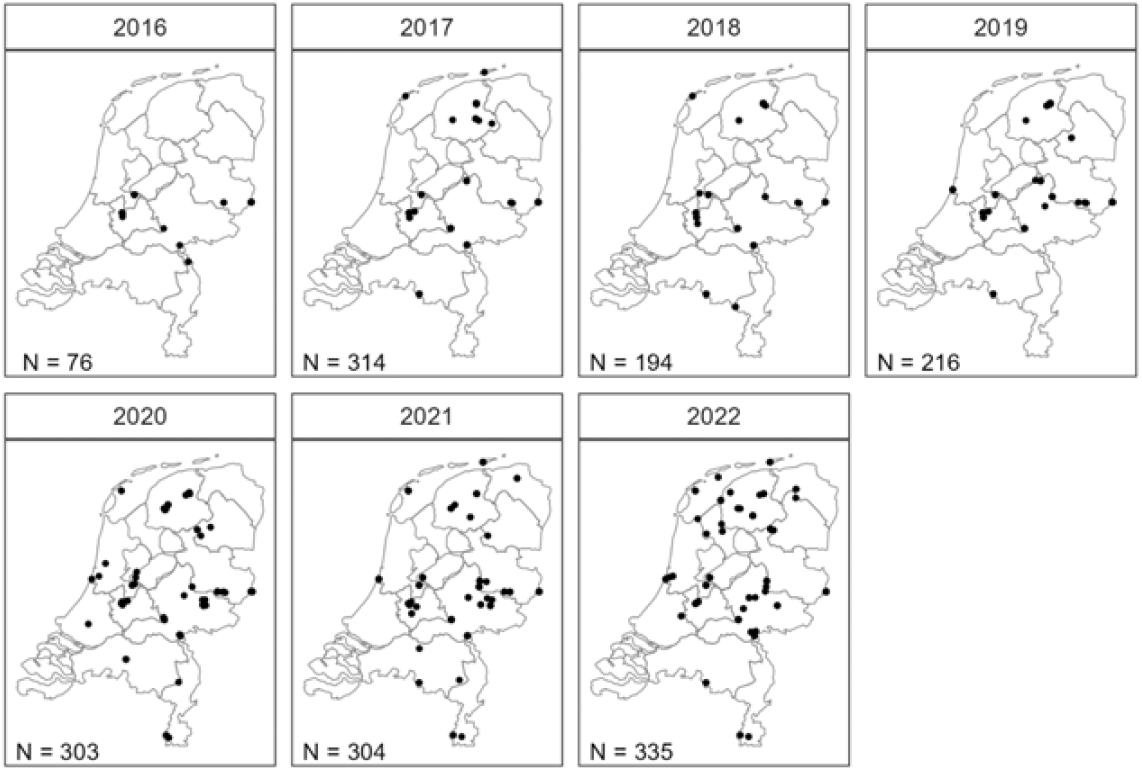
Annual maps of sampling locations, where each dot represents one sample.

### Supplement: Identifiability analysis

We evaluated the ability of our model to estimate parameters from simulated data. We created three simulated datasets using three different sets of parameters. The number of individuals in each simulated dataset was equal to that in the real dataset (N=1742). For each simulated dataset, we evaluated the median and 95% credible interval of the posterior distributions and compared this to the input parameters.

**Supplementary figure 2.**
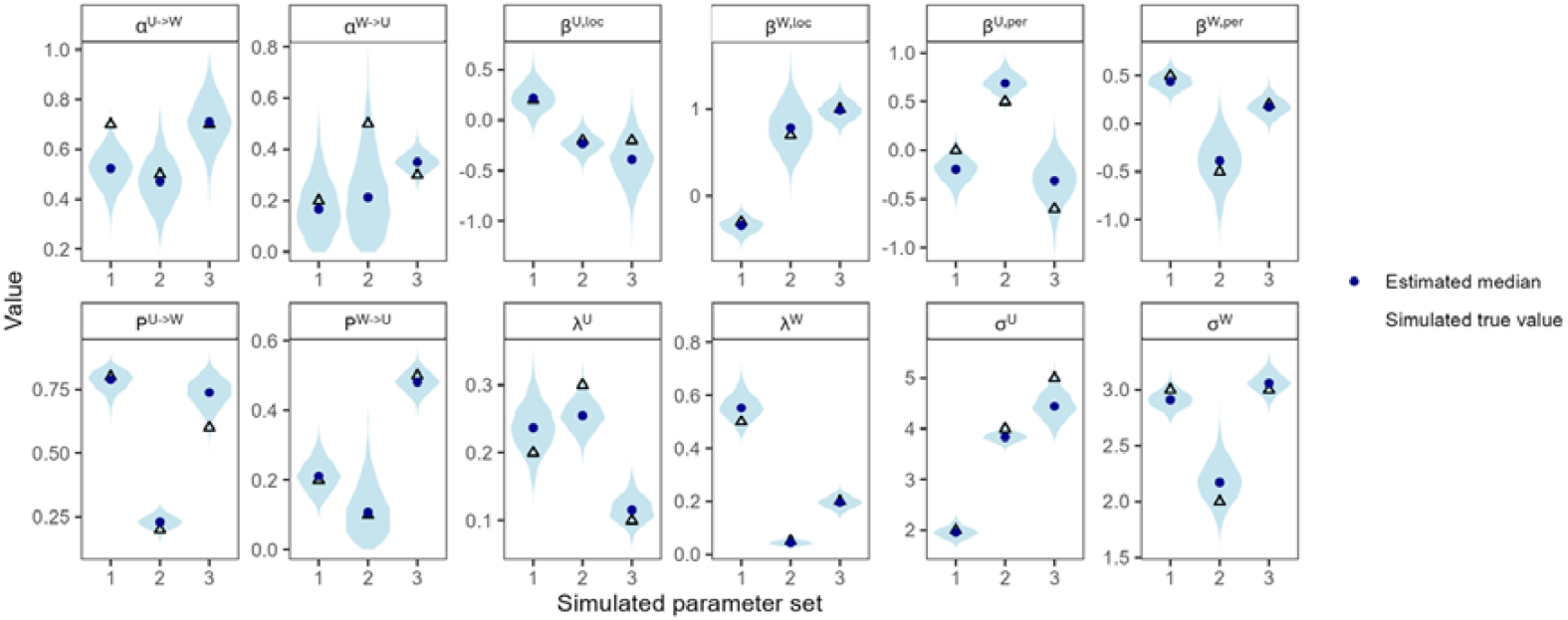
Estimated parameters from three simulated datasets (1-3) with the true value used in the simulation (blue triangle), the estimated median (blue circles) and posterior distributions (blue violins). Each estimate was based on three independent chains with 2000 accepted particles each.

### Supplement: Diagnostics

**Traceplots**

**Supplementary figure 3.**
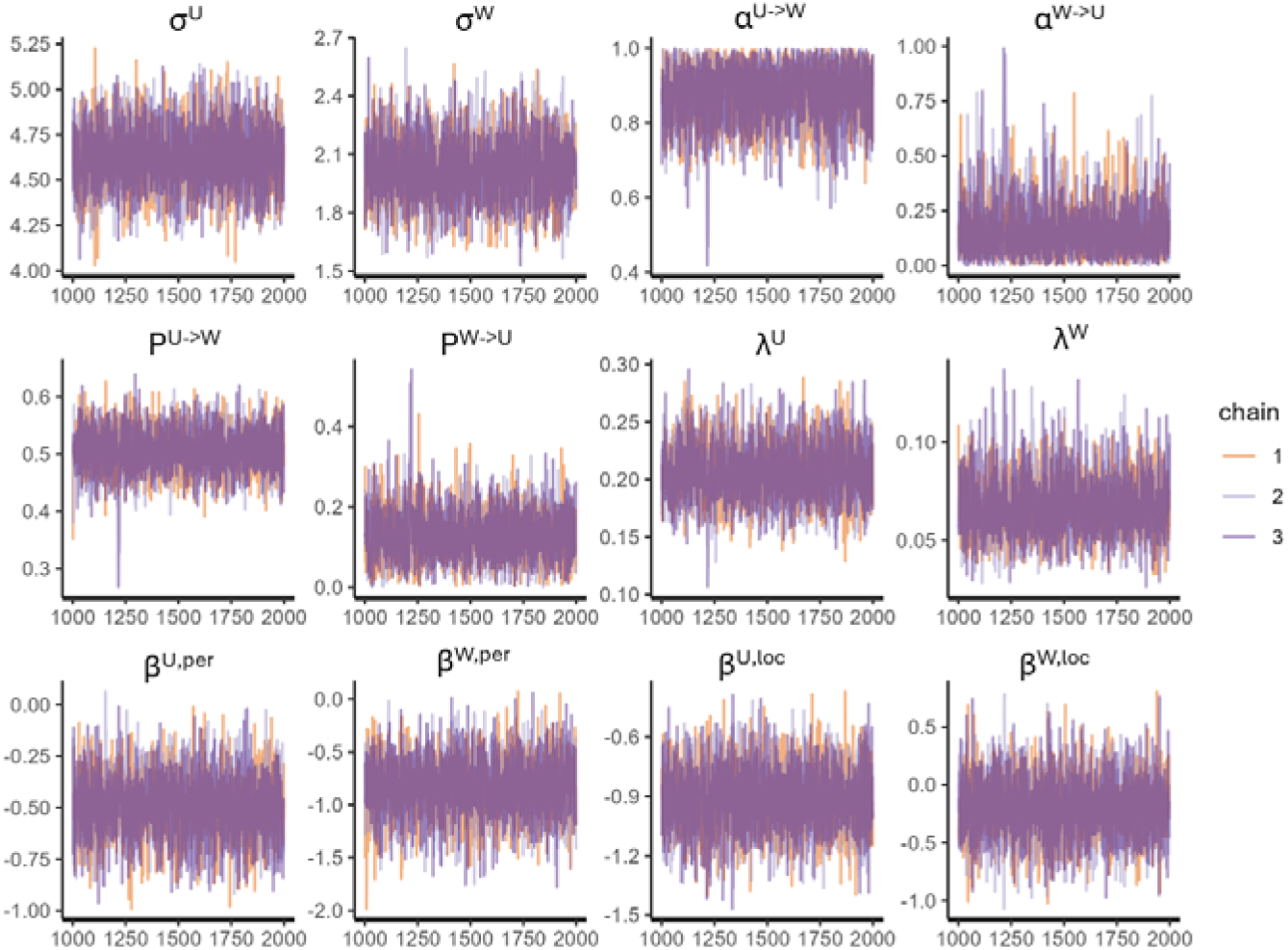
Trace plots

### Supplement: Posterior distributions and model fit

**Supplementary figure 4.**
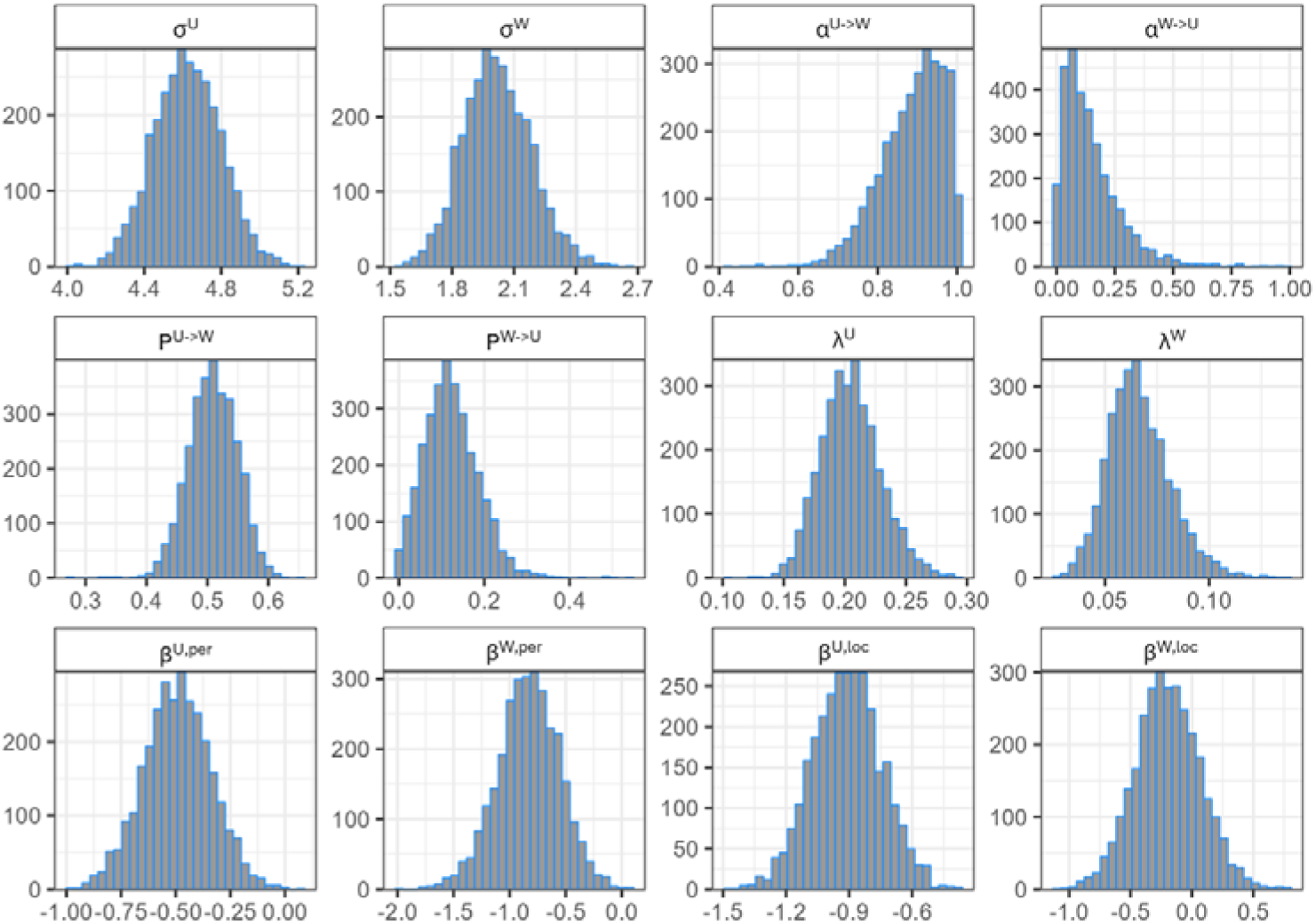
Posterior distributions of estimated parameters

#### Model fit to data for full model

**Supplementary figure 5:**
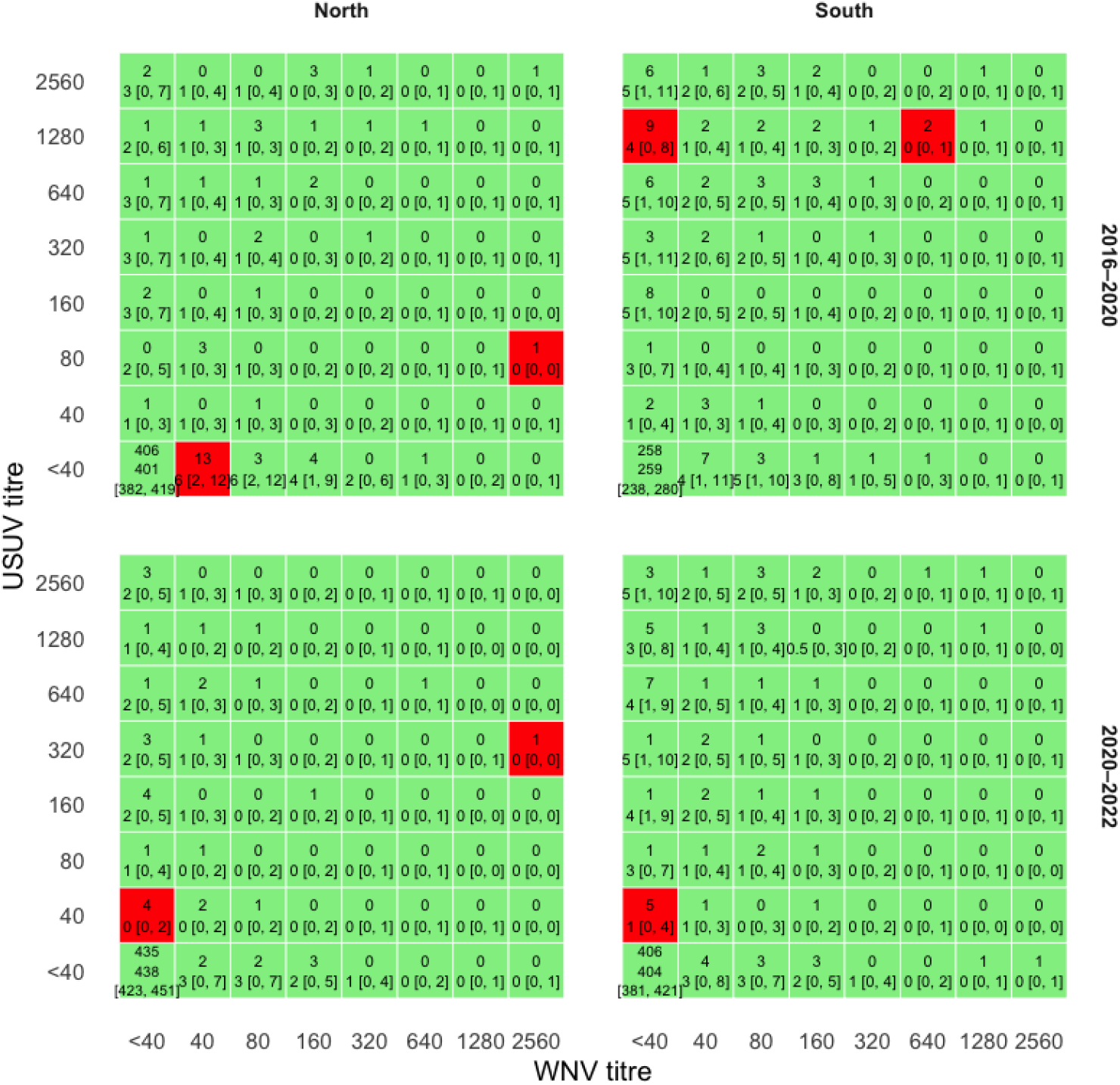
Agreement of simulated vs observed titre frequency for full model: frequency table of titre combinations by period and region. Each cell shows the number of samples in the observed data (top value in each cell) and the simulated data (bottom value for the median and 95% confidence interval). Cells are coloured green where the true value falls within the range and red otherwise.

#### Model fit to data for model with seroprevalence fixed at uncorrected estimates from current sero-classification algorithm

**Supplementary figure 6:**
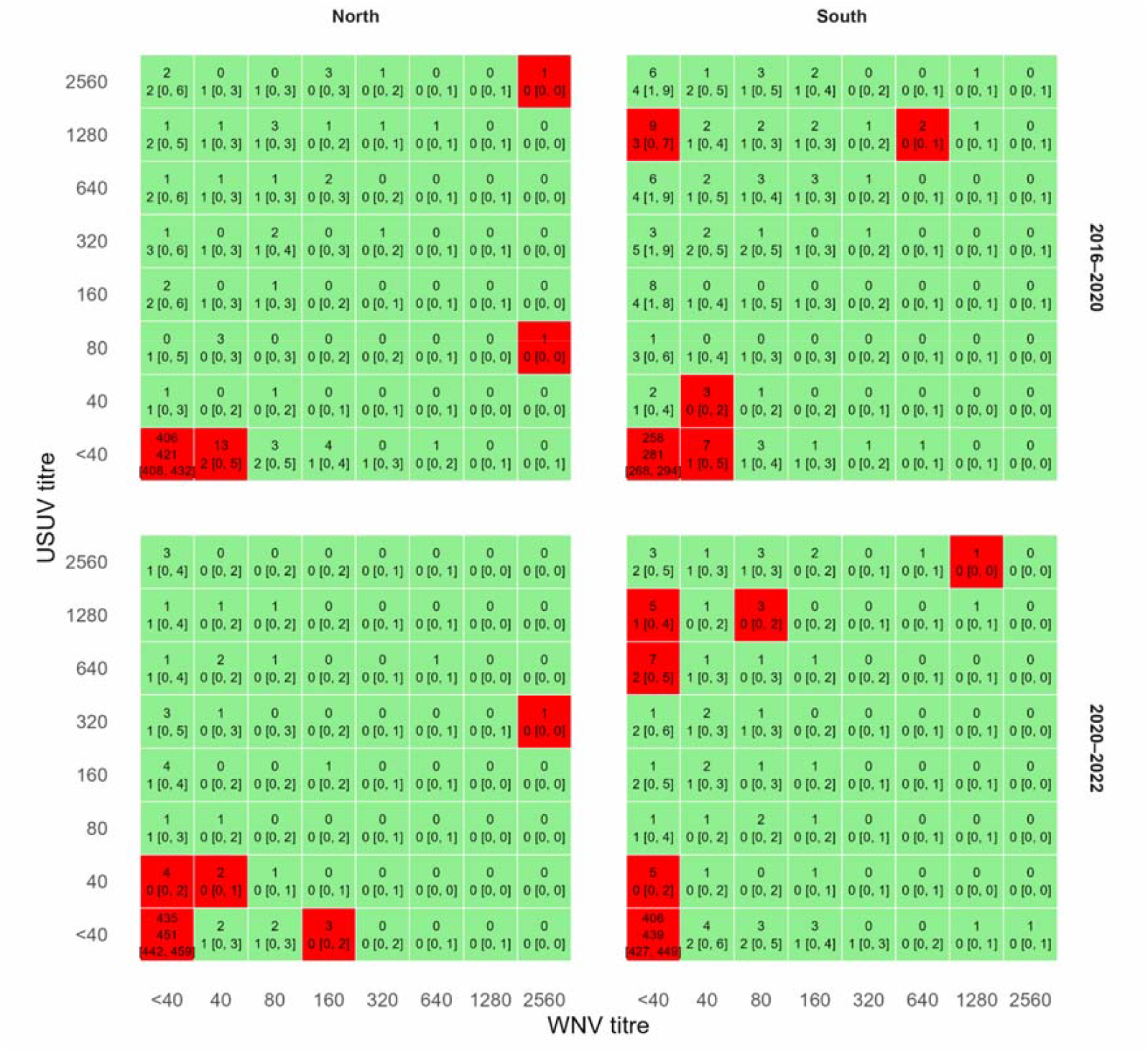
Agreement of simulated vs observed titre frequency for model with seroprevalence fixed at uncorrected estimates from current sero-classification algorithm: frequency table of titre combinations by period and region. Each cell shows the number of samples in the observed data (top value in each cell) and the simulated data (bottom value for the median and 95% confidence interval). Cells are coloured green where the true value falls within the range and red otherwise.

### Supplement: Impact of infection-induced mortality on parameter estimates

In Supplementary Figure 7A, we represent the relationship between the true attack rate and seroprevalence for various values of the infection mortality rate. The value for infection mortality risk used in our analysis was obtained from de Wit et al. [9] (median 74%, 95%CI 66-78).

In this study we only reported the seroprevalence and attack rate, but it is important to note that the infection mortality and the discrepancy between the seroprevalence and attack rate also has consequences for the estimates of the relative FOI across different regions 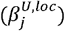 and periods 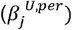. If *p*_1_ =1−*e* ^−λ^and *p*_2_ =1−*e* ^−λ × *β*^ are the estimated infection probabilities, then the relative force of infection *β* based on the true attack rates *q*_l_ and *q*_2_ given by formula (10) above, is given by

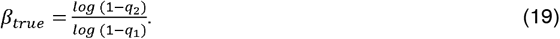

We represent in Supplementary Figure 7B the relationship between *β*_true_ and *β* for different values of the model-based seroprevalence.

**Supplementary Figure 7.**
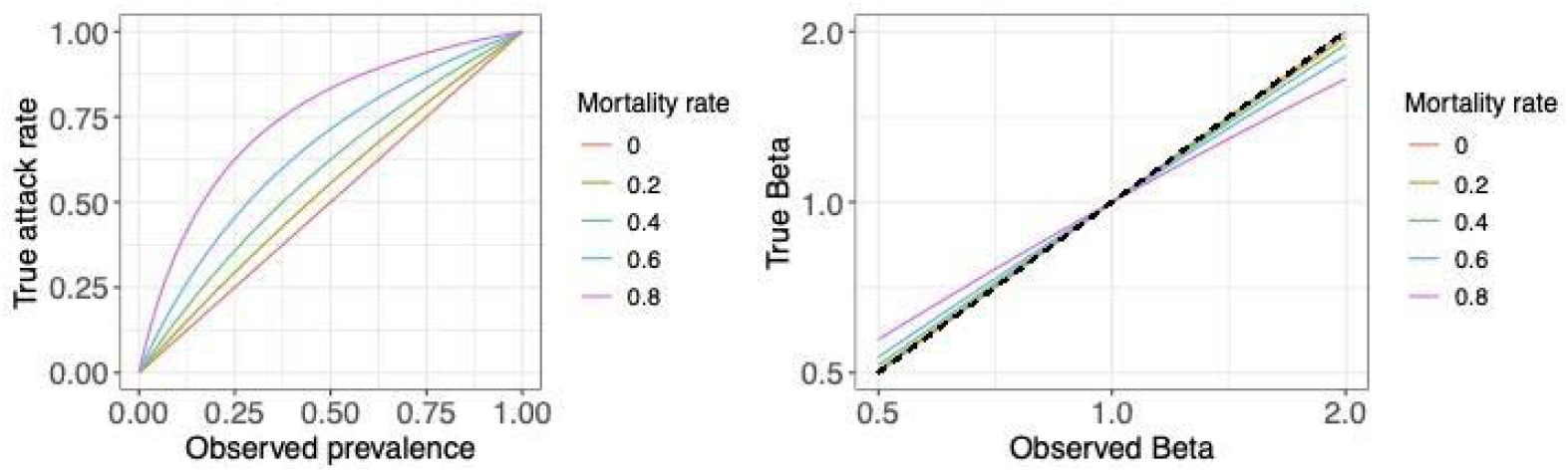
Correction of the seroprevalence for a potentially lethal infection. **(A)** Comparison of the true attack rate and the observed seroprevalence (eq. (17)), for different values of the infection mortality rate. **(B)** Relationship between the true and observed relative infection probabilities (*β*) across groups for different mortality rates eq (19).

Additionally, we investigated whether infection-induced mortality and fluctuating circulation levels influenced the estimation of the attack rate in the context of co-circulation. To explore this, we simulated a scenario where two viruses circulated over multiple periods. At the end of each period, a survey was conducted, with the data stratified into two age groups: young and old birds. Old birds are young birds of the previous period and therefore may have been exposed during both periods. Additionally, natural mortality was incorporated into the model.

We ran simulations across various scenarios (Supplementary figure 8) to assess whether the relationship between observed seroprevalence and the true attack rate holds under these more complex conditions. We found that the attack rate could be accurately recovered from the observed seroprevalence using the same correction formula eq. (16)-(17) as in the single-virus case, provided that mortality rates were known. This indicates that the correction approach remains valid in the presence of co-circulation, multiple periods, and additional sources of mortality.

**Supplementary Figure 8.**
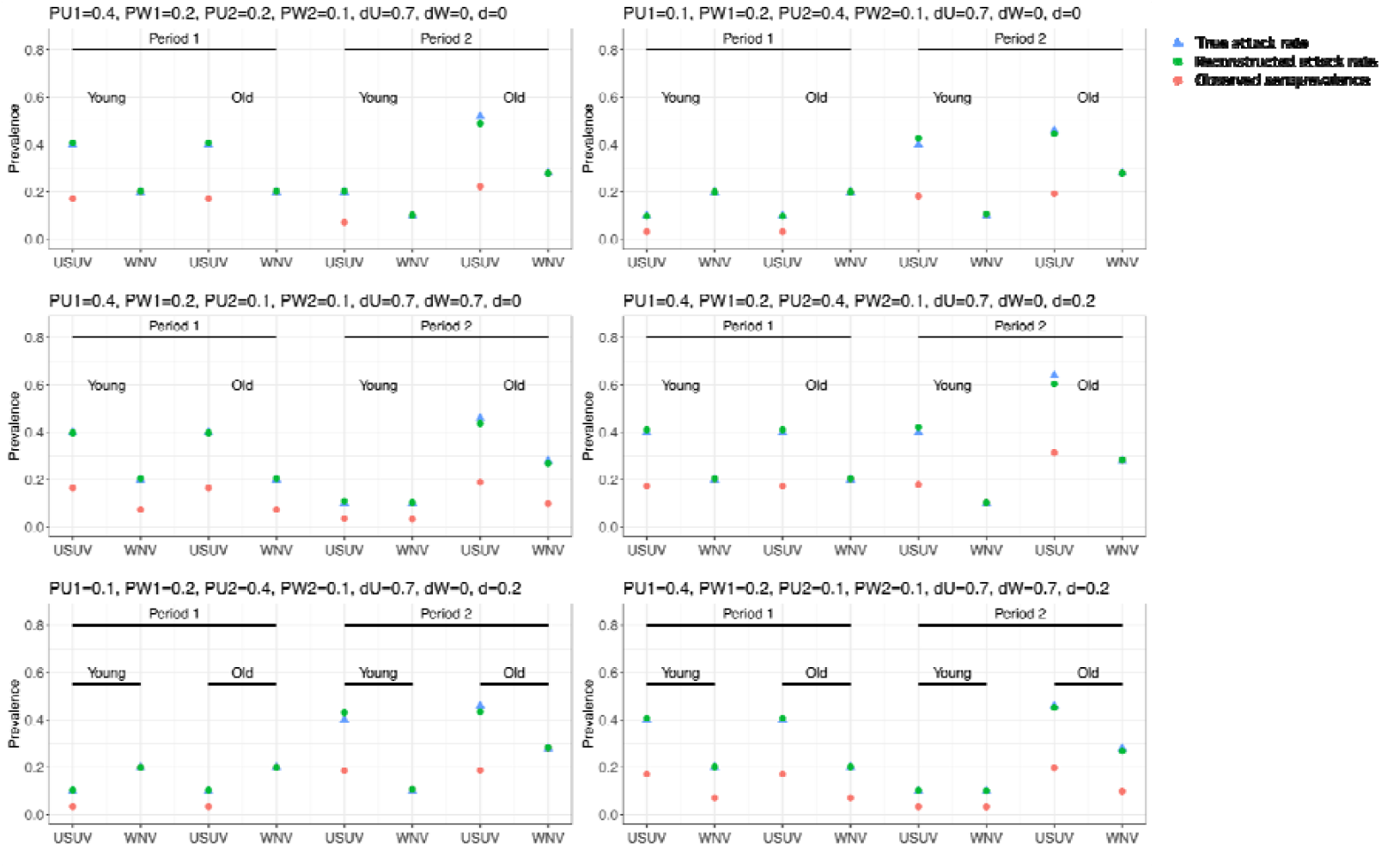
Attack rate estimation in simulations. Infections of young and old birds were simulated over different periods and for different scenarios, corresponding to various attack rates on each period (PU1, PU2, PW1, PW2), infection-induced mortality (dU and dW), natural mortality rate over each period (d). Colours indicate the input attack rate (blue), observed (red) and reconstructed (green).

### Supplement: Titre classification

#### Details of calculation sensitivity and specificity

A short description of how sensitivity and specificity were calculated was described in the methods. Here, we provide a more detailed explanation for WNV, the same logic was applied to calculate the sensitivity and specificity for USUV.

The true positive rate was calculated by summing across all titre combinations classified as WNV-positive the probability of this titre multiplied by the probability that this titre was due to WNV.

The false negative rate was calculated by summing across all titre combinations classified as WNV-negative the probability of this titre multiplied by the probability that this titre was due to WNV.

The true negative rate was calculated by summing across all titre combinations classified as WNV-negative the probability of this titre multiplied by the probability that this titre was not due to WNV.

The false positive rate was calculated by summing across all titre combinations classified as WNV-positive the probability of this titre multiplied by the probability that this titre was not due to WNV.

## Notes

### Competing Interest Statement

The authors have declared no competing interest.

